# Analysis of CheW-like domains provides insights into organization of prokaryotic chemotaxis systems

**DOI:** 10.1101/2022.05.07.491037

**Authors:** Luke R. Vass, Robert B. Bourret, Clay A. Foster

**Affiliations:** Department of Microbiology & Immunology, University of North Carolina, Chapel Hill, North Carolina, United States of America

**Keywords:** Bacterial chemotaxis systems, CheW-like domains, CheA, CheW, CheV

## Abstract

The ability to control locomotion in a dynamic environment provides a competitive advantage for microorganisms, thus driving the evolution of sophisticated regulatory systems. Nineteen known categories of chemotaxis systems control motility mediated by flagella or Type IV pili, plus other cellular functions. A key feature that distinguishes chemotaxis systems from generic two-component regulatory systems is separation of receptor and kinase functions into distinct proteins, linked by CheW scaffold proteins. This arrangement allows for formation of varied arrays with remarkable signaling properties. We recently analyzed sequences of CheW-like domains found in CheA kinases and CheW and CheV scaffold proteins. Sixteen *Architectures* of CheA, CheW, and CheV proteins contain ∼94% of all CheW-like domains and form six *Classes* with likely functional specializations.

We surveyed chemotaxis system categories and proteins containing CheW-like domains in ∼1900 prokaryotic species, the most comprehensive analysis to date, revealing new insights. Co-occurrence analyses suggested that many chemotaxis systems occur in non-random combinations within species, implying synergy or antagonism. Furthermore, many *Architectures* of proteins containing CheW-like domains occurred predominantly with specific categories of chemotaxis systems, suggesting specialized functional interactions. We propose *Class* 1 (∼80%) and *Class* 2 (∼20%) CheW proteins exhibit preferences for distinct chemoreceptor structures. Furthermore, rare (∼1%) Class 6 CheW proteins frequently co-occurred with methyl-accepting coiled coil (MAC) proteins, which contain both receptor and kinase functions and so do not require connection via a CheW scaffold but may benefit from arrays. Lastly, rare multi-domain CheW proteins may interact with different receptors than single domain CheW proteins.

## 1 INTRODUCTION

Two component signaling systems are found in bacteria, archaea, and certain eukaryotes such as plants and fungi.^1, 2^ Two-component pathways allow organisms to sense and respond to environmental stimuli in an organized and timely manner (reviewed in ^2^). The most basic two-component system consists of a membrane-bound sensor histidine kinase that binds ATP and modulates an autophosphorylation reaction in response to an external stimulus. The resulting phosphoryl group is transferred to an aspartate residue in the receiver domain of a downstream response regulator to elicit an appropriate cellular response. However, two-component pathways often exhibit more complexity, incorporating additional proteins to form branching signaling networks (reviewed in ^3^). This increased complexity provides an opportunity to fine-tune the signal-response characteristics of a given pathway, tailoring the system to better suit the needs of the organism (various advantages are summarized in ^4^).

The chemotaxis pathway is one of the most well-studied two-component systems^5^ and is present in some form in nearly every motile microorganism. Chemotaxis is a regulatory strategy used to direct the movement of an organism towards resources (attractants) or away from undesirable substances (repellants). Variations on the chemotaxis system allow for locomotion in response to a variety of physicochemical parameters such as temperature, pH, magnetism, etc., in addition to nutrients.^6–10^ The pathway utilizes a diverse repertoire of transmembrane environmental sensors to detect properties of interest. The sensors, known as chemoreceptors (also called methyl-accepting chemotaxis proteins, or MCPs), typically form mixed transmembrane arrays with remarkable, highly customizable signaling properties, including wide dynamic ranges, integration of mixed inputs, cooperativity, and rapid signal amplification potential.^11^ The sophisticated information processing capabilities of the chemotaxis system are advantageous not only for general survival, but also for invasion-, colonization-, and virulence-related processes in pathogenic microorganisms.^12–16^

The chemotaxis pathway of *Escherichia coli* has been thoroughly characterized and is an example of a two-component system that incorporates additional proteins to achieve a more rapid and coordinated response.^17^ The pathway begins at a transmembrane array of chemoreceptors. Activation of the sensor array depends on detection of an environmental stimulus and the methylation status of the receptors. Following detection, a stimulus is converted into receptor conformational changes to initiate processing and propagation. The signal is then passed to the histidine kinase, CheA, which integrates information from multiple receptors through autophosphorylation after summing positive and negative stimuli.^18, 19^ The signal path then splits into “excitation” and “adaptation” branches. In the excitation path, phosphoryl groups are passed to the response regulator CheY. Phosphorylation alters the equilibria between active and inactive conformations in the CheY population, which ultimately modulates flagellar motor behavior and motility. The adaptation path in *E. coli* features CheR and CheB. CheR includes a methyltransferase domain that steadily adds methyl groups to the chemoreceptors, independent of environmental stimuli. CheB is a response regulator that includes a methylesterase domain whose activity is tightly regulated by phosphorylation and removes methyl groups from the chemoreceptors in response to a sufficient environmental change. The adaptation path forms a delayed negative feedback loop, imparting a “memory” to the system and allowing the organism to both follow a stimulus gradient and reset upon reaching a uniform environment. Some chemotaxis systems also incorporate separate phosphatases, such as CheZ, to catalyze the removal of phosphoryl groups and terminate the response at a specific point in the pathway.^20^ Many organisms encode multiple chemotaxis systems for regulating multiple forms of propulsion and/or gene expression^5^.

An important distinction between a generic two-component pathway and a chemotaxis system is the separation of sensor and kinase functions into distinct protein species. Physical separation allows CheA kinases to integrate information from many different chemoreceptors, substantially enhancing the utility of the system.^18, 19, 21^ This integration is facilitated by CheW proteins, which act as scaffolds between the various receptors and CheA kinases.^22, 23^ The classical architecture of CheA includes a histidine phosphotransfer (Hpt) domain (Pfam ID PF01627) containing the site of phosphorylation (a His residue), a dimerization domain (PF02895), an HATPase_c ATP binding and catalytic domain (PF02518), and a CheW-like domain (PF01584).^24^ The CheW-like domain in *E. coli* CheA interacts with its counterpart CheW-like domain in standalone CheW proteins and also with cytoplasmic portions of the receptors to form MCP signaling arrays. The formation of supramolecular MCP•CheW•CheA oligomers is an essential part of the system and leads to CheA activation, signal propagation, and ultimately a downstream shift in flagellar behavior and/or locomotion.^21, 23, 25^

Most existing information on CheW-like domains describes the canonical, standalone CheW protein (the most abundant form in nature).^25^ The interactions between the free species of CheW and its partners (MCPs and CheAs) have been well-characterized for several microbial species, particularly *E. coli.*^26–30^

The most common occurrence of CheW-like domains other than CheA- or CheW-lineage proteins (-lineage referring to proteins that can be characterized as analogous to the canonical CheA and standalone CheW proteins in *E. coli*) is fused to a receiver domain in CheV proteins (reviewed in ^22, 31^). Various commonly studied organisms, including *Bacillus subtilis*, *Helicobacter pylori,* and *Vibrio cholerae,* encode one or more CheV proteins. Although CheV proteins are present in approximately one-third of all chemotaxis systems, their role(s) are still poorly understood.^32^ CheV is thought to be involved in both CheA modulation/MCP adaptation and array formation/polar localization.^33, 34^ The receiver domain of CheV may also serve as a general phosphate sink for the system.^31, 35^

A landmark study by Wuichet and Zhulin established an evolutionary classification of chemotaxis signaling systems in prokaryotes^32^, summarized here. The core components of essentially all chemotaxis systems, as outlined above, are the MCP•CheW•CheA arrays, the CheB and CheR adaptation enzymes, and CheY response regulators. Chemotaxis systems often have multiple MCPs and CheW proteins, and it is technically difficult to distinguish CheY from other single domain response regulators. Therefore, to provide a consistent foundation, the original classification was based on phylogenetic trees of CheA/CheB/CheR sequences and supported by certain other phylogenetic markers. There are 19 known categories of standard chemotaxis systems, each containing characteristic arrangements of *che* genes, distinct sets of auxiliary components (CheC phosphatase, CheD deamidase, CheV scaffold protein, CheX phosphatase, and/or CheZ phosphatase), and often unique architectures for certain core components. Seventeen categories of chemotaxis systems (F1 through F17) control flagellar motility, one controls Type IV pili (Tfp), and one controls alternative (non-motility) cellular functions (ACF). In addition, there are two categories of related chemotaxis systems based on methyl-accepting coiled coil (MAC) proteins, which contain both receptor and kinase functions, rather than separate MCP and CheA proteins.

A recent companion paper describes our classification of CheW-like domains into six *Classes*, likely related to specific functional specializations.^36^ Nearly all (∼94%) CheW-like domains are encompassed by 16 distinct *Architectures* (Figure 1). CheW-like domains in proteins with the CheW.I *Architecture* belong to three different *Classes*, whereas CheW-like domains in other *Contexts* predominantly correspond to single *Classes*. Most CheW-like domains in CheW- and CheV-lineage proteins belong to *Class* 1. Most CheW-like domains in CheA-lineage proteins belong to *Class* 3, except for the CheA.VII and CheA.X *Architectures* (*Class* 4, which contain multiple Hpt domains) and the *Class* 5 C-terminal CheW-like domains in CheA proteins with two such domains. Rare (∼1%) *Class* 2 CheW-like domains are found in CheW.I *Architectures* and exhibit properties of both CheA-lineage and CheW-lineage proteins. About 20% of CheW-like domains in CheW-lineage proteins (*Class* 6) appear subtly different from *Class* 1 and may form yet another specialized *Class*. Strikingly, this study provides insights into both Class 2 and Class 6 CheW-like domains.

**FIGURE 1.**
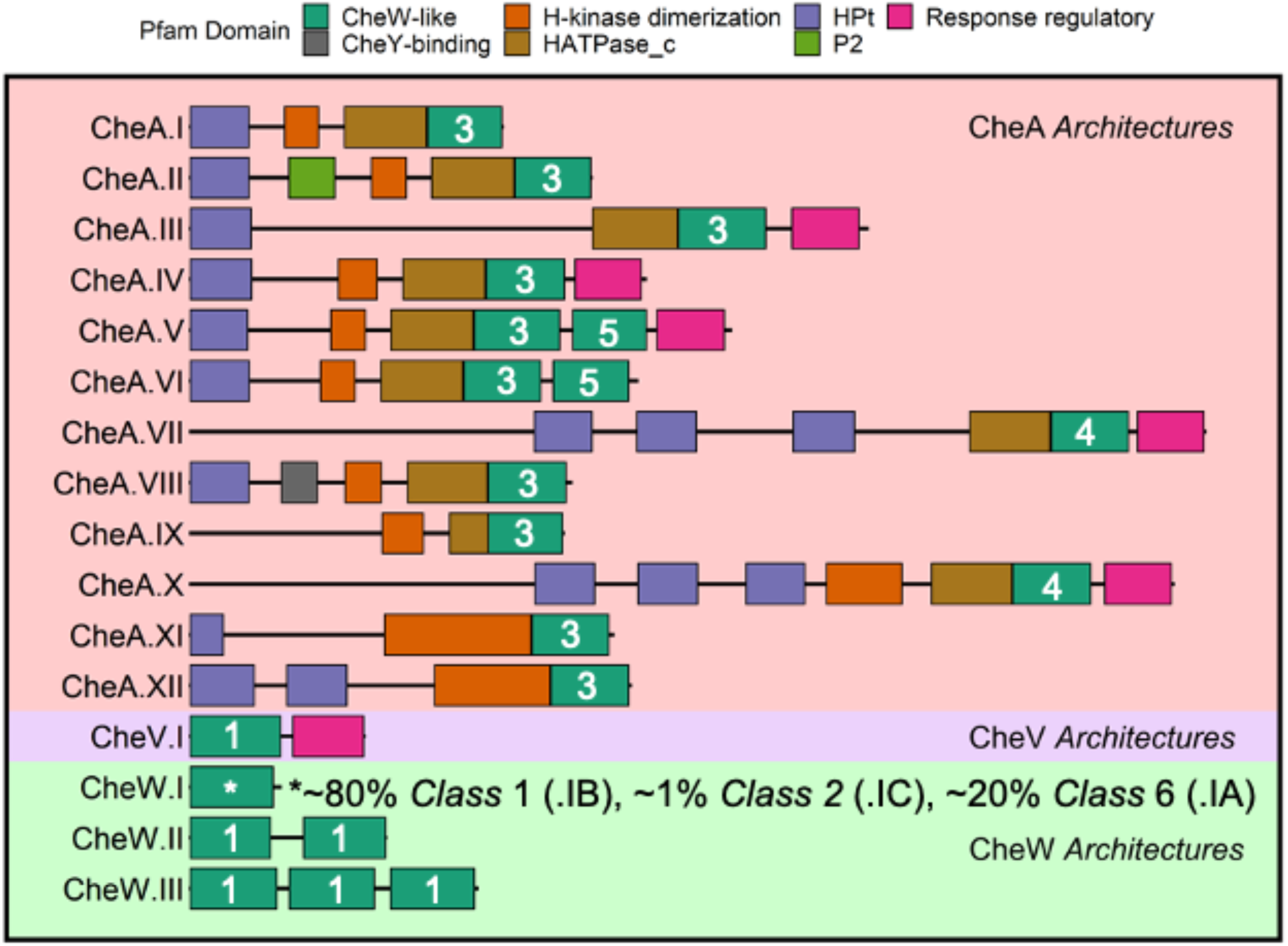
Major *Architectures* of proteins that contain CheW-like domains. (from ^36^). CheA-lineage *Architectures* are designated by a Roman numeral suffix in order of decreasing abundance. CheW-lineage *Architectures* are designated by a Roman numeral suffix indicating the number of CheW-like domains. The CheW.I *Architecture* includes sequences that belong to three distinct *Classes*, designated CheW.IA, CheW.IB, and CheW.IC. Classes of CheW-like domains are indicated by white numerals.

In this work, we combined our classification scheme of CheA-, CheW-, and CheV-lineage (CheW-like domain containing) proteins^36^ with that of Wuichet and Zhulin for other chemotaxis proteins^32^ to gain additional insights into the occurrence and organization of chemotaxis systems. We found that (i) many chemotaxis system categories occur in non-random combinations within microbial species, and (ii) specific *Architectures* of CheA/CheW proteins are preferentially associated with specific chemotaxis system categories, suggesting functional interactions.

## 2 MATERIALS AND METHODS

### 2.1 Protein sequence database and co-occurrence analysis

Analysis of CheW-like domains (PF01584) was described in ^36^, using sequences sourced from Representative Proteome 35 (RP35)^37^ and the Pfam database (version 33, obtained May 2020).^38^ We previously extracted all proteins that contained CheW-like domains and belonged to the 16 *Architectures* (Figure 1) that account for ∼94% of CheW-like domains within the RP35 dataset.^36^

To analyze the co-occurrence (presence-absence) patterns of the various chemotaxis components encoded by proteomes within the same dataset, proteins containing one or more of the following domains were extracted: MCP (PF00015); CheR (PF01739); CheB (PF01339); CheD (PF03975); CheZ (PF04344); CheCX (also simply called CheC, PF04509/PF13690).^38^ Receiver domain-containing proteins orthologous to CheY were excluded for the sake of interpretability. Full protein sequences (excluding the previously analyzed CheW-containing proteins) were scanned and classified with HMMER3 (version 3.3) using the previously described chemotaxis system models (utilized by the MiST database; version 3.0).^32, 39–41^ Thus, MCPs were classified by number of heptad repeats,^42^ whereas CheB, CheC (including closely related CheX^43^), CheD, CheR, and CheZ proteins were assigned to the chemotaxis system categories of Wuichet and Zhulin.^32^ A combined frequency table with organisms in columns and chemotaxis components in rows was generated by merging the previously classified CheW-containing *Architectures* with the other chemotaxis components by organism. Components with fewer than 20 positive occurrences were discarded from the analysis. The excluded components belonged to eight categories (F3 or F11 through F17) of chemotaxis systems, which were also previously observed to be infrequent.^32^ The resulting count matrix of distinct chemotaxis components encoded by 1887 distinct proteomes (Dataset S1) was used to generate a heatmap (using the R package ComplexHeatmap, version 2.7.11) of the co-occurrence patterns within the representative proteomes.^44^ The taxize package in R (version 0.9.99.947)^45^ was used to assign putative phyla and classes to the batch of relevant organisms (sourced from the NCBI Taxonomy Browser^46^). Assignments were visualized as row/column annotations with ComplexHeatmap. Rows (components) and columns (species) were grouped in an unsupervised manner by hierarchical clustering, based on Spearman correlation, using Ward’s clustering method (option “ward.D2”). The resulting dendrograms (both row and column) were split by height (using the cutree() function, as implemented by the row_split/column_split options) into putative functional “blocs” of chemotaxis components. The choice of height does not meaningfully affect the order of rows and columns in the heatmap but does affect the number of blocs/splits displayed. We selected a single height to cut the row dendogram such that components from the same chemotaxis system categories largely ended up in the same row bloc (i.e. all F1 components together, all F6 components together, etc.), making strong biological sense. The height to cut the column dendogram was chosen to optimize interpretability, i.e., visual segregation of data into discrete column blocks

To analyze the co-occurrence patterns of the chemotaxis classes themselves, rather than the individual components, the original matrix (Dataset S1) generated to create Figure 2 was binarized. Rows featuring CheA, CheW, CheV and MCP paralogs were first removed because they are used in multiple chemotaxis system categories. A binary scheme was then applied for each individual chemotaxis system category (F1, F2, F4, F5, F6, F7, F7.z, F8, F9, F10, Tfp, ACF, MAC1, MAC2 and Uncat). If a given organism contained at least one of the corresponding components (CheB/C/D/R/Z) for a chemotaxis category, then the category was considered present in the final table (=1), irrespective of the presence/absence of other chemotaxis system categories (or paralogous instances of the same category). Those categories lacking all components were assigned absent (=0). A small percentage of organisms (<5%) lacked auxiliary components entirely and were excluded. Dataset S2 provides a full breakdown with corresponding proteome counts. A new heatmap (Figure 3) was generated in a similar manner using ComplexHeatmap to re-cluster the proteomes (i.e., columns; 1797 distinct organisms). Row order was maintained to match Figure 2. Dataset S3 provides the relative abundances of the various chemotaxis system categories across all proteomes.

**FIGURE 2.**
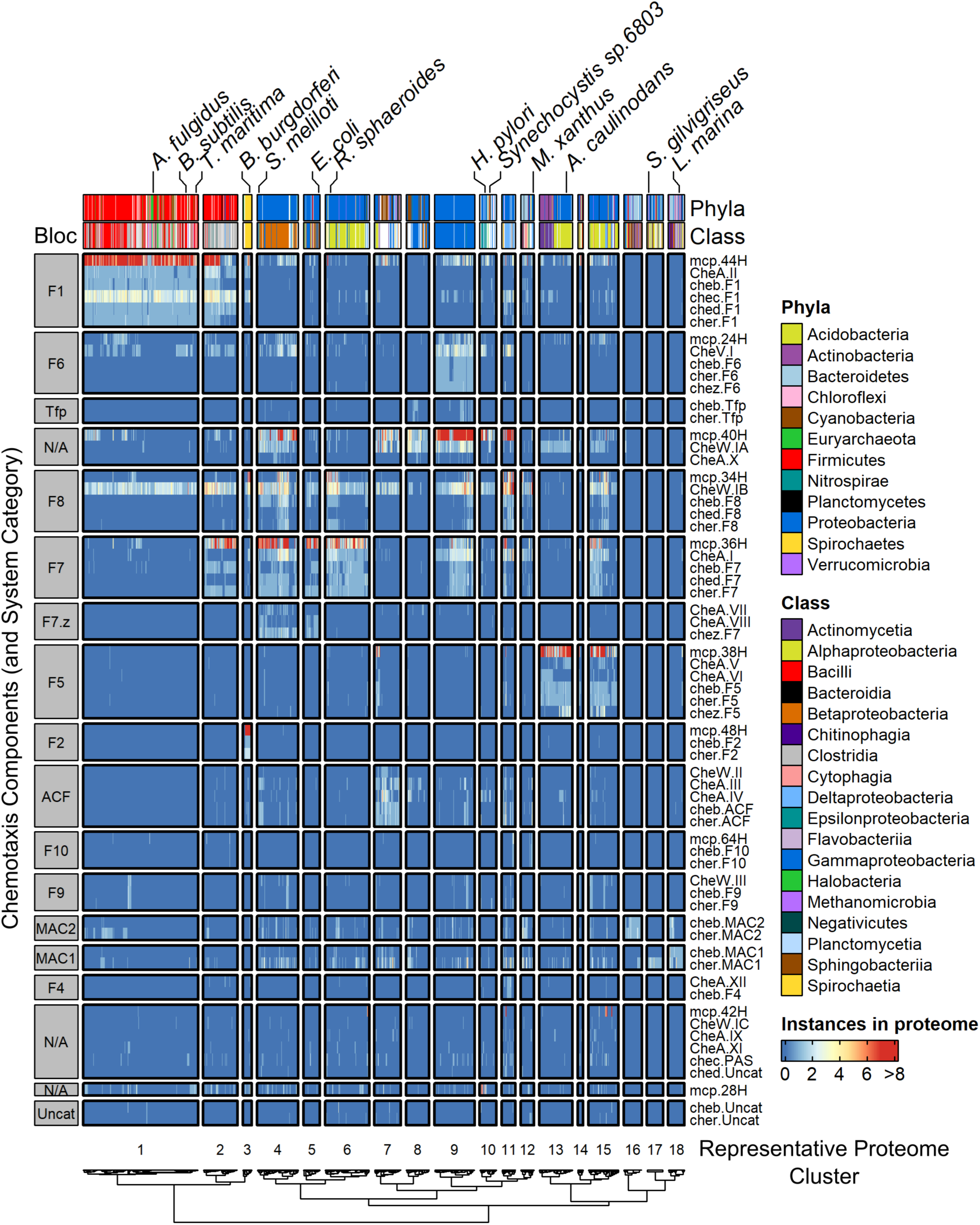
Co-occurrences of individual chemotaxis components in RP35 representative proteome set. Total occurrence counts were used. Components with < 20 occurrences were excluded. Column annotations were shaded by Phyla (groups with < 10 occurrences are unlabeled) and Class (groups with < 10 occurrences are unlabeled). Notable organisms were tagged. Results were split by dendrogram height into functional “blocs” by clustering both proteomes (columns) and chemotaxis components (rows). Representative Proteome clusters were labeled as 1-18, whereas component blocs were labeled with most likely chemotaxis system category. Row dendogram shown in Figure S1.

**FIGURE 3.**
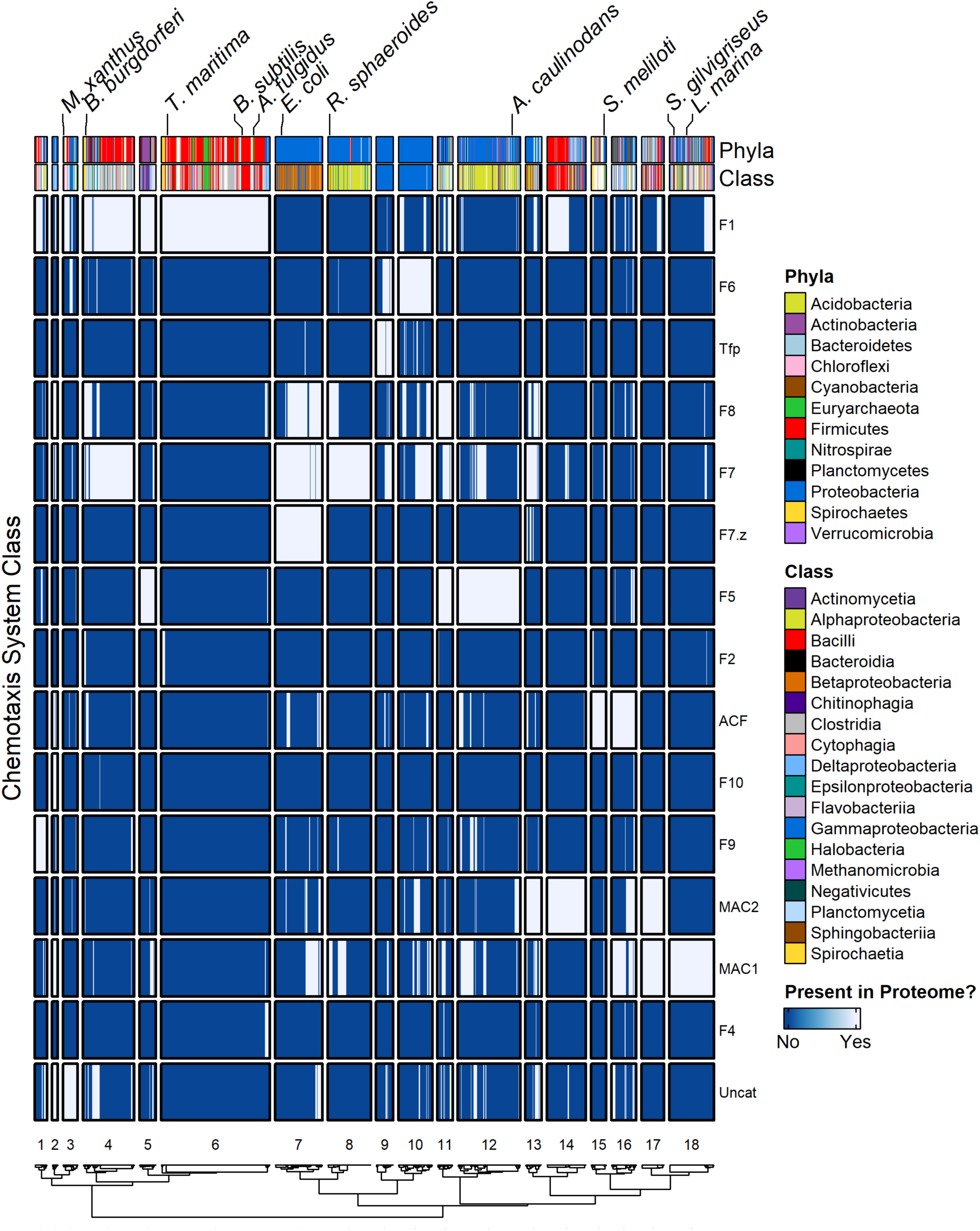
Simplified co-occurrence schematic of chemotaxis system categories in RP35 representative proteome set. A binary presence/absence scheme was used for visualization. A chemotaxis system category was determined to be present in a given proteome if at least one of the following components was detected of the appropriate category: CheB/C/D/R/Z. CheA/V/W and MCP components were excluded from the analysis, because some of these components function with more than one chemotaxis system category. Notable organisms were labelled. Row order was maintained for consistency with Figure 2, meaning categories were not clustered. Results were split by dendrogram height into functional “blocs” by clustering proteomes (columns). However, because the datasets upon which Figures 2 and 3 are based are different, the resulting proteome clusters and cluster numbers are different than in Figure 2.

### 2.2 Phyletic direct coupling analysis (PhyDCA) and network generation

The traditional approach to phylogenetic profiling utilizes some form of local correlation metric, such as Hamming distance or Pearson correlation, to transform a simple binary matrix representing presence (1) or absence (0) of specific components/proteins/genes in various species into a corresponding interaction network. However, classical profiling suffers from several disadvantages, such as the influence of “intermediate” effects on apparent direct couplings (meaning that if A co-evolves with B, and B co-evolves with C, A may also appear to co-evolve with C). A more recent approach introduced the concept of direct coupling analysis, a statistical modeling technique able to distinguish between direct and more indirect co-evolutionary signals, to the profiling of presence-absence patterns.^47^ This method, called Phyletic Direct Coupling Analysis, or PhyDCA, has demonstrated substantially increased accuracy compared to the more traditional correlation-based approaches and provided a convenient means by which to quantify the relationships presented in Figure 2. The frequencies used to create the Figure 2 co-occurrence heatmap (covering the full, non-filtered complement of chemotaxis components) were converted into a binary phylogenetic profile matrix to analyze pairwise evolutionary couplings. Data were analyzed with PhyDCA (using the mfDCA implementation) to estimate relevant quantitative phyletic pairings using a global statistical modelling approach.^47^ The phyletic coupling (*J_i,j_*) between two domains and/or components in our data was used to estimate the favorability of finding multiple elements within the same species, corresponding to the principle that a biological process (i.e., chemotaxis) would require both components to function and produce a strong positive coupling. A negative coupling could also be interpreted as alternative solutions for similar functionality in a given system. We took the 125 (4%) strongest positive pairwise couplings (i.e., the presence of one component favors the presence of the other) and created a non-directed graph to visualize the web of co-evolutionary signals using the R packages igraph and ggraph (using the Fruchterman-Reingold algorithm).^48–50^ The phyletic coupling scores of the pairs displayed in Figure 4 (≥ 0.44) were well within the range of known significantly predictive scores for the PhyDCA method (as determined previously by comparing with known pairwise relations as a reference set).^47^ The complete list of pairwise coupling strengths (>3,000 possible pairs) is in Dataset S4.

**FIGURE 4.**
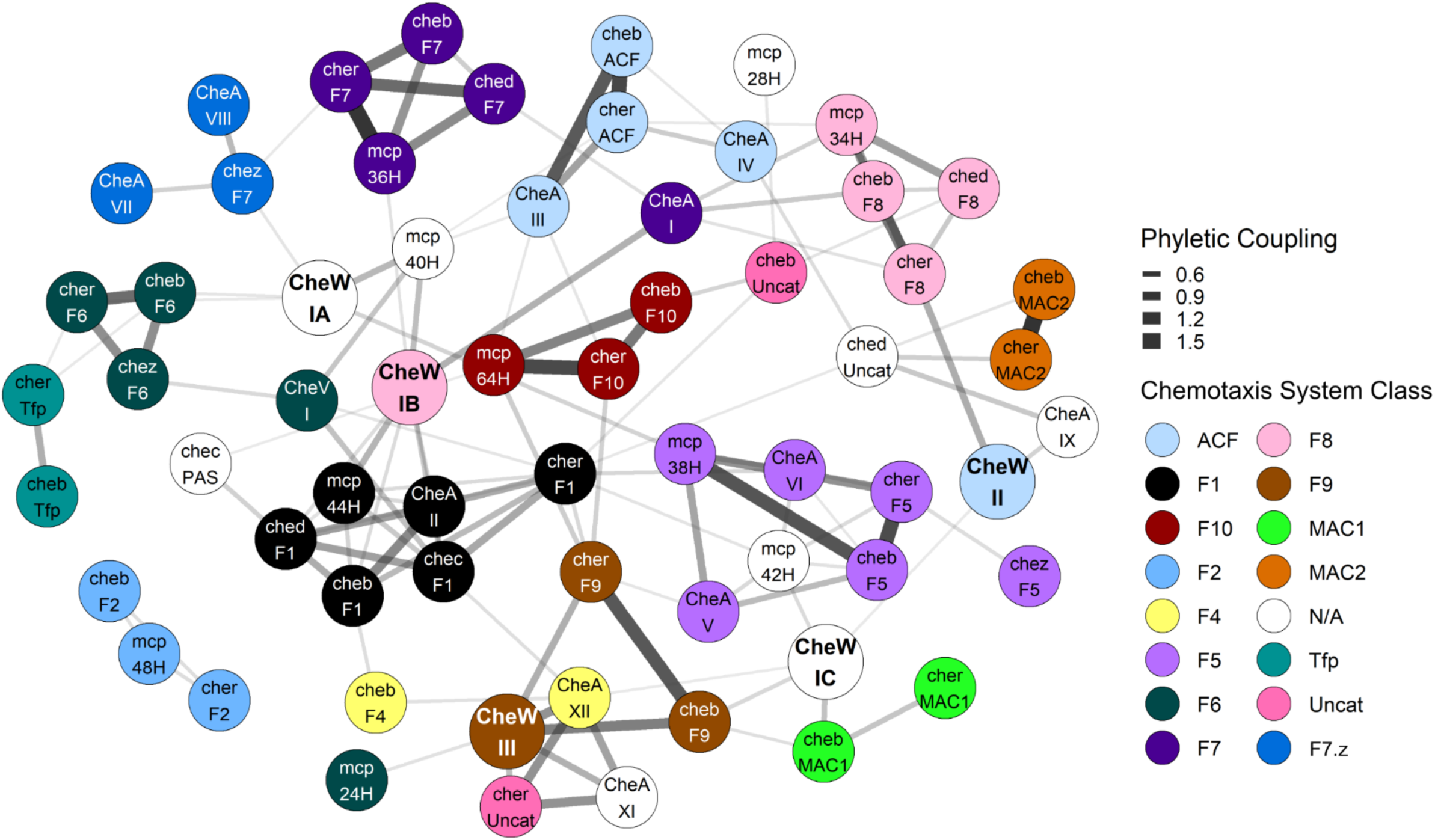
Network representation of inferred phyletic couplings between *Architectures* containing CheW-like domains and remaining chemotaxis system components. Co-occurrence data of chemotaxis components extracted from the RP35 representative proteome set were converted to a binary phylogenetic profile matrix. Phyletic Direct Coupling Analysis (PhyDCA) was used to quantify the favorability (correlation) of chemotaxis components co-occurring within the same organism. Strong favorability/high coupling typically corresponds to a cellular function (i.e., chemotaxis) requiring both components, though not necessarily to a direct biophysical interaction. The top 125 (∼4%) positive co-evolutionary pairings were used to construct a graph based on phyletic coupling strength. Architectural assignments correspond to those included in Figure 1 (i.e., identical thresholds). Edge width was scaled with phyletic coupling strength. Notes: *Architecture* CheA.X did not appear in the top 4% strongest phyletic couplings and was excluded from the graph. cheb.F2, cher.F2, and mcp.48H formed a cluster disconnected from the rest of the network, but all three coupled to CheA.II at slightly lower strengths (top ∼10%) (Dataset S4).

## 3 RESULTS

### 3.1 Many combinations of chemotaxis system categories are non-randomly distributed across prokaryotic species

To better understand the organization of prokaryotic chemotaxis signal transduction pathways, we first extracted all identifiable chemotaxis proteins except CheYs from 1887 prokaryotic proteomes as described in Materials and Methods. Each type of chemotaxis protein (CheA, CheB, etc.) was sorted as follows: MCPs were classified by the number of heptad repeats,^42^ proteins containing CheW-like domains were classified by *Architecture* as in Figure 1, and all other chemotaxis proteins were assigned to the chemotaxis system categories of Wuichet and Zhulin.^32^ The end result was 64 distinct chemotaxis components (Dataset S1). We then organized the information in two dimensions, with species in columns and chemotaxis components in rows. Hierarchical clustering was performed to optimally group organisms and chemotaxis components with similar co-occurrence patterns. The results were visualized in a composite heatmap, with color indicating the number of instances of each component in a proteome (Figure 2).

Wuichet and Zhulin classified chemotaxis system categories primarily based on phylogenetic trees of CheA, CheB, and CheR protein sequences.^32^ The dominant feature of Figure 2 is that when clustered in an unsupervised manner the occurrence (not sequence) data primarily formed recognizable blocs as a function of putative chemotaxis system category (row dendrogram shown in Figure S1) and proteome. Such a phenomenon strongly suggested that the data were linked in both dimensions, across chemotaxis system categories and across proteomes. If each species (proteome) encoded a single category of chemotaxis system, then two-dimensional clustering would be trivial, and proteomes would group perfectly into functional blocs by system category. However, >50% of all prokaryotic genomes that encode chemotaxis systems contain multiple systems (first determined in ^32^ and again corroborated by our work). With many known categories of chemotaxis systems, if combinations featuring multiple categories in a single species were random, then clustering by species would be disrupted, precluding the previously described functional blocs. Therefore, we infer that many of the naturally occurring combinations of chemotaxis system categories with prokaryotic species are non-random. We explore in qualitative and quantitative detail multiple aspects of the relationships between chemotaxis components, chemotaxis system categories, and proteomes in the following sections.

### 3.2 Presence-absence analysis reveals preferred and disfavored category combinations in organisms with multiple chemotaxis systems

By converting the frequency table of components (Dataset S1) used to generate Figure 2 into a simplified, binary presence/absence matrix featuring only the chemotaxis system categories themselves, we next determined the most common naturally occurring combinations of chemotaxis system categories (Dataset S2) and generated a heatmap displaying the presence/absence of each chemotaxis system category across species (Figure 3).

We first sought to compare the distribution of chemotaxis system categories across species with the evolutionary relationships between the chemotaxis systems. The phylogenetic tree of chemotaxis systems features three main branches (here arbitrarily designated Branches 1, 2 and 3), with Branch 2 exhibiting three sub-Branches.^32^ The most common 10% of chemotaxis category combinations observed in Dataset S2 accounted for two-thirds of the proteomes in our study and are displayed in relation to the various Branches in Table 1, with cross-referencing to their locations in Figure 3. Approximately a third of the proteomes encoded only a single category of chemotaxis system (Table 1, top).

**TABLE 1.**
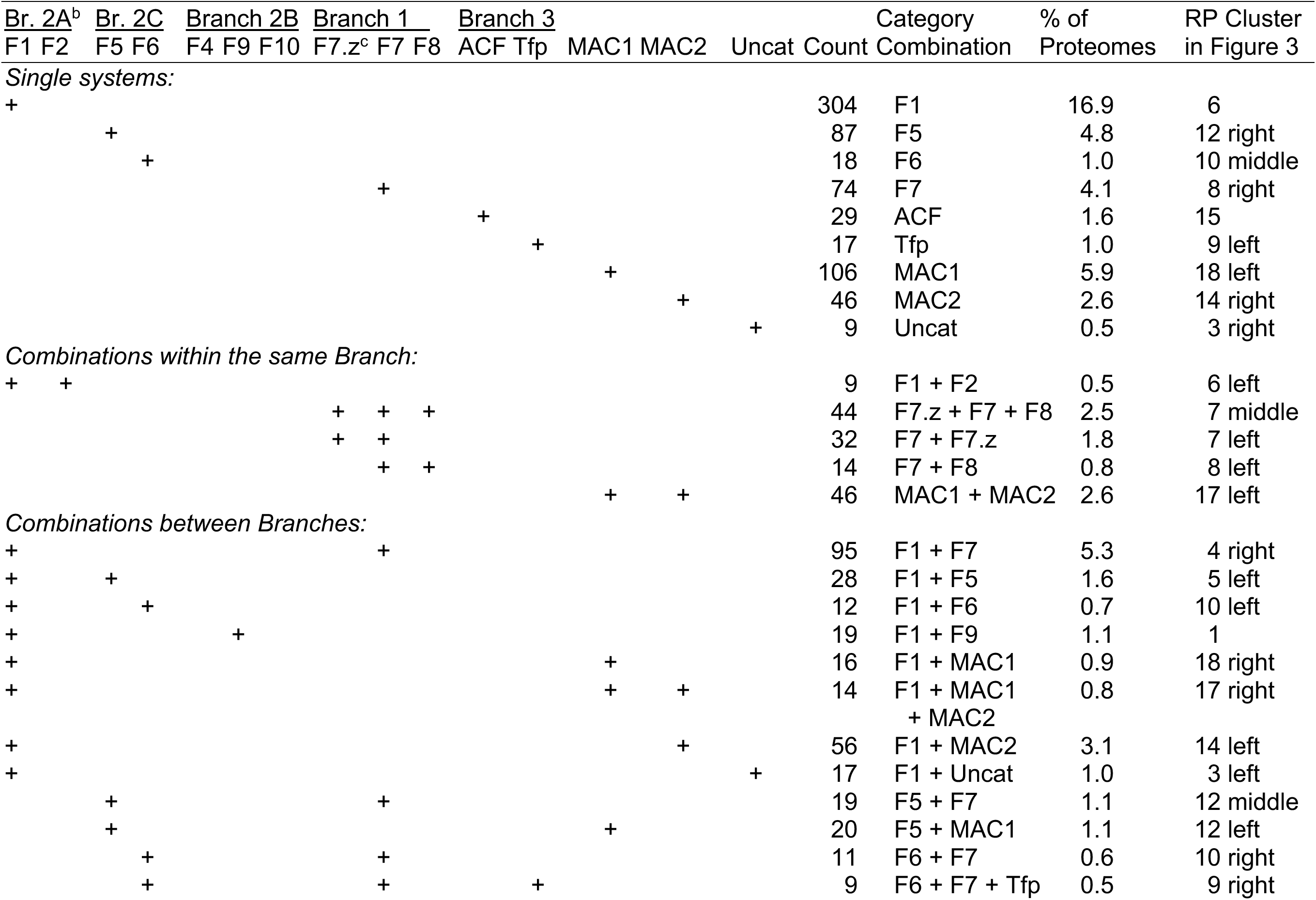
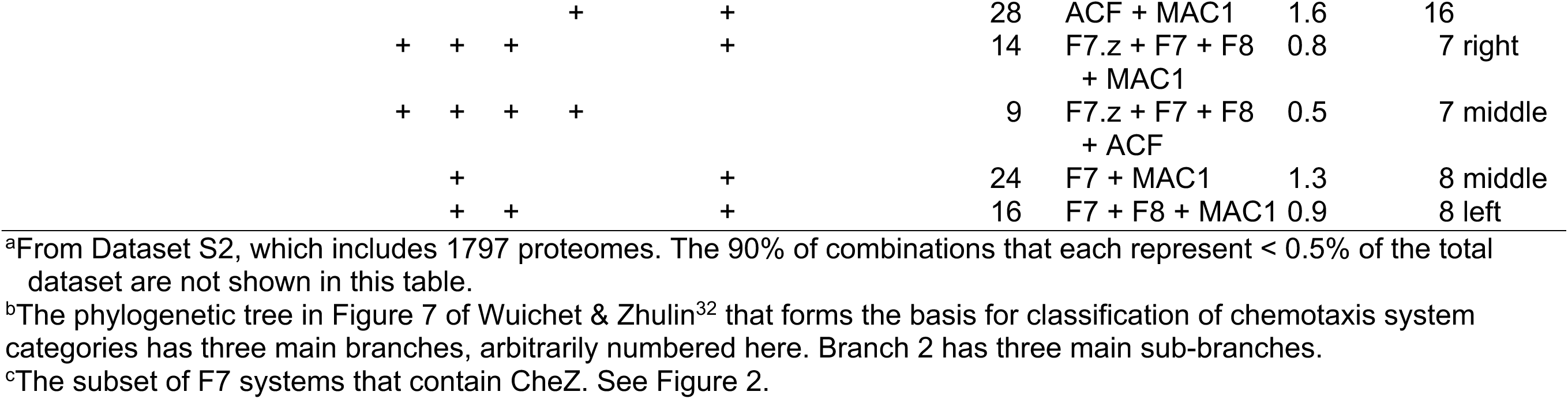
Most common chemotaxis system category combinations in Representative Proteomes^a^.

We next focused on the proteomes encoding multiple chemotaxis systems, first examining pairwise combinations of categories found within the same Branch (Table 1, middle). In general, flagellar chemotaxis system categories within the same Branch (F1/F2, F4/F9/F10, and F7.z/F7/F8) co-occurred at substantially higher frequencies in Dataset S2 than would be expected based on frequencies of the constituent categories from Dataset S3 (see calculations in Table S1). Conversely, the combination of categories F5/F6 occurred approximately five times less frequently than expected using the same relationship, implying some form of negative selection. It seems plausible that the components of the two categories (F5/F6) may interfere with one another (or share overlapping functions to some degree). In contrast to the flagellar systems, categories ACF/Tfp and MAC1/2 both co-occurred at frequencies consistent with a random distribution.

Over 50% of all proteomes encoding multiple distinct chemotaxis systems in our dataset included categories from disparate Branches of the classification tree (Table 1 bottom and Dataset S2), suggesting highly diverse origins for most systems. We found that most outgroup combinations occurred at frequencies relatively consistent with a random distribution (Table S2). A notable exception was class F7.z, which occurred with categories in outgroup Branches much less frequently than expected (consistent with a strong/semi-exclusive linkage between F7.z/F7 and F7.z/F8). Additionally, class F9 co-occurred with both F5 and F8 systems more frequently than expected from a random distribution.

Finally, we examined the co-occurrent relationships between the flagellar chemotaxis system classes and the ACF/Tfp/MAC1/MAC2 systems. *A priori*, we speculated that the non-flagellar systems would operate independently from the flagellar-controlling classes, revealing no obvious selective pressure(s). While the majority of pairwise combinations between flagellar and non-flagellar systems (approximately 67%) supported our prediction, a full third deviated substantially (Table S3). One-quarter of pairwise combinations were observed at lower-than-expected frequencies, with nearly half of the cases of negative selection involving either F1 or F2 systems. Ten percent of pairwise combinations were observed at higher-than-expected frequencies, with nearly half of the cases of positive selection involving F10 systems.

### 3.3 Matching *Architectures* of proteins containing CheW-like domains to preferred chemotaxis system categories

Examining the functional blocs revealed by clustering in Figure 2 suggested that various *Architectures* of CheA and CheW proteins are differentially favored by divergent chemotaxis systems. In particular, many *Architectures* clustered with components belonging to one chemotaxis system category. Wuichet and Zhulin described eight cases of distinct architectures for proteins containing CheW-like domains that were characteristic of specific chemotaxis system categories.^32^ The four *Architectures* noted by Wuichet and Zhulin that were sufficiently abundant to be included in our study were CheA.III/CheA.IV, CheA.VI, CheA.XII, and CheW.III, which were linked to categories ACF/F3, F5, F4, and F9, respectively. Our data confirmed most previous assignments. Our larger sample size enabled us to also propose multiple additional assignments. Qualitative assignments of specific *Architectures* to specific chemotaxis systems inferred from sorting patterns in Figure 2 are summarized in Table 2. Quantitative analyses reported in Section 3.4 strengthen and extend these observations. The 17 new relationships identified in our work link *Architectures* CheA.I, CheA.II, CheA.V, CheA.VII, CheA.VIII, CheV.I, CheW.IA, CheW.IB, CheW.IC, and CheW.II with chemotaxis system categories F7/F8, F1, F5, F7.z, F7.z, F1/F6, F5/F6/F7.z, F1/F7/F8, MAC1, and F8/ACF, respectively.

**TABLE 2.**
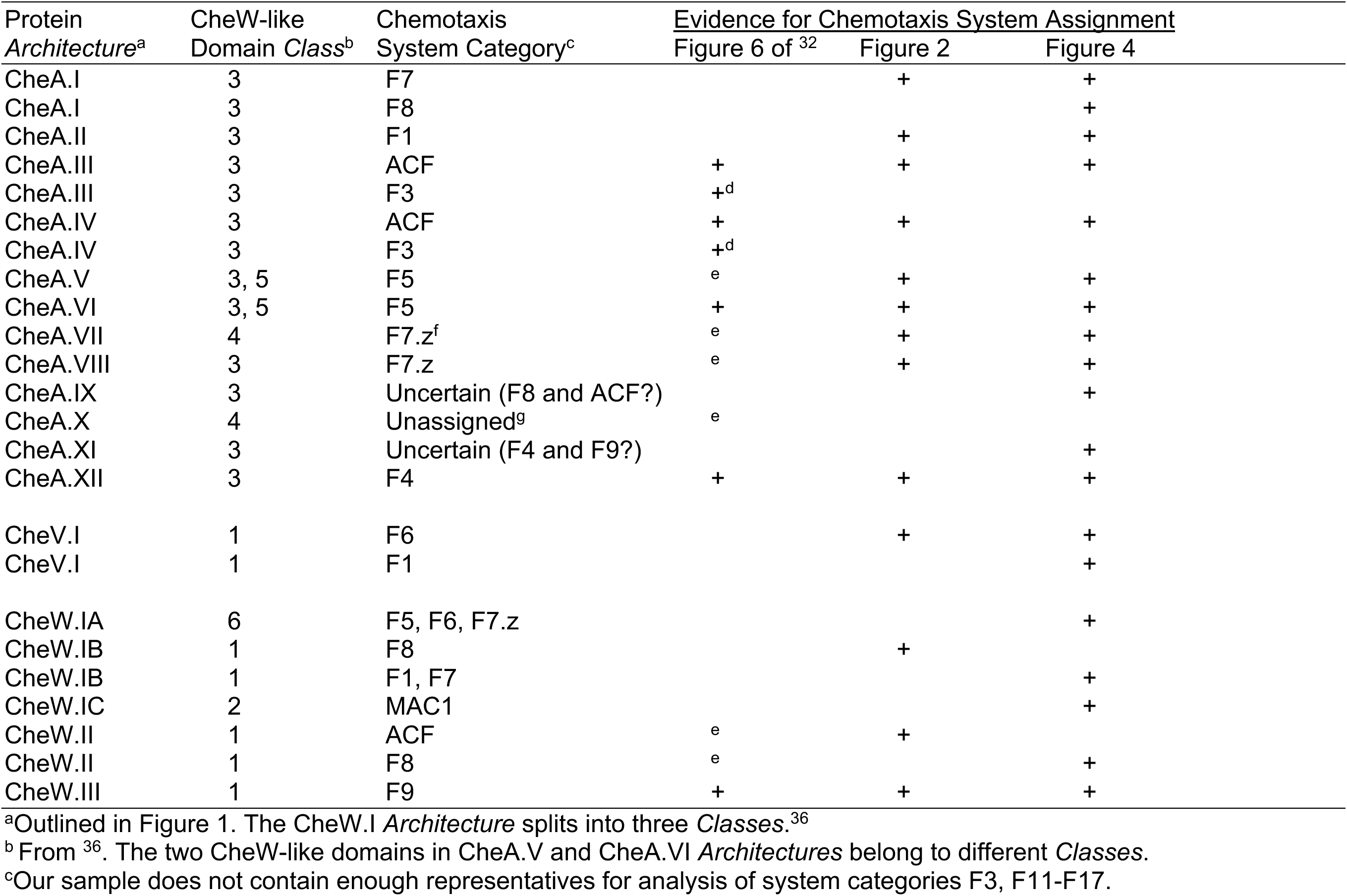
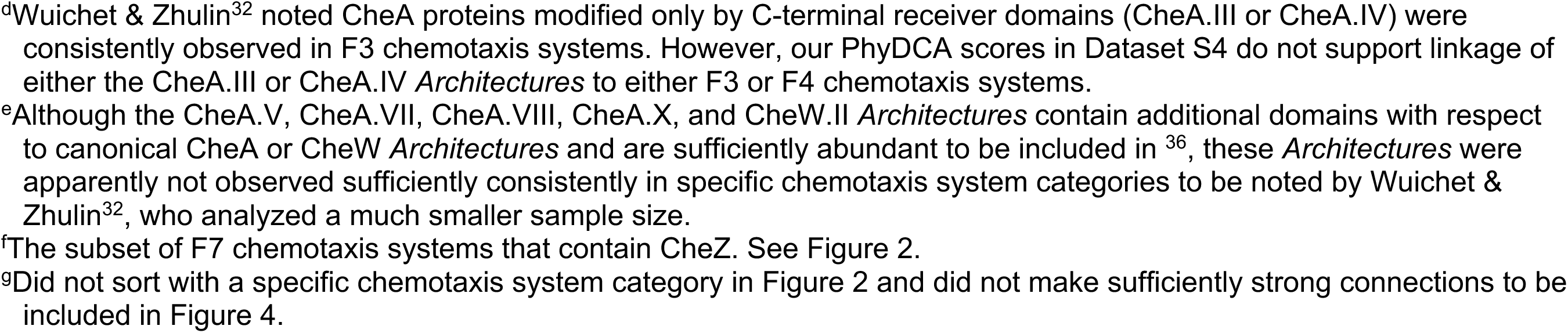
Assignment of CheA and CheW protein *Architectures* to chemotaxis system categories.

| 812 | Protein | CheW-like | Chemotaxis | Evidence for Chemotaxis System Assignment |  |  |
| --- | --- | --- | --- | --- | --- | --- |
| 813 | *Architecture*<sup>a</sup> | Domain *Class*<sup>b</sup> | System Category<sup>c</sup> | Figure 6 of <sup>32</sup> | Figure 2 | Figure 4 |
| 814 | CheA.I | 3 | F7 |  | + | + |
| 815 | CheA.I | 3 | F8 |  |  | + |
| 816 | CheA.II | 3 | F1 |  | + | + |
| 817 | CheA.III | 3 | ACF | + | + | + |
| 818 | CheA.III | 3 | F3 | +<sup>d</sup> |  |  |
| 819 | CheA.IV | 3 | ACF | + | + | + |
| 820 | CheA.IV | 3 | F3 | +<sup>d</sup> |  |  |
| 821 | CheA.V | 3, 5 | F5 | e | + | + |
| 822 | CheA.VI | 3, 5 | F5 | + | + | + |
| 823 | CheA.VII | 4 | F7.z<sup>f</sup> | e | + | + |
| 824 | CheA.VIII | 3 | F7.z | e | + | + |
| 825 | CheA.IX | 3 | Uncertain (F8 and ACF?) |  |  | + |
| 826 | CheA.X | 4 | Unassigned<sup>g</sup> | e |  |  |
| 827 | CheA.XI | 3 | Uncertain (F4 and F9?) |  |  | + |
| 828 | CheA.XII | 3 | F4 | + | + | + |
| 829 |  |  |  |  |  |  |
| 830 | CheV.I | 1 | F6 |  | + | + |
| 831 | CheV.I | 1 | F1 |  |  | + |
| 832 |  |  |  |  |  |  |
| 833 | CheW.IA | 6 | F5, F6, F7.z |  |  | + |
| 834 | CheW.IB | 1 | F8 |  | + |  |
| 835 | CheW.IB | 1 | F1, F7 |  |  | + |
| 836 | CheW.IC | 2 | MAC1 |  |  | + |
| 837 | CheW.II | 1 | ACF | e | + |  |
| 838 | CheW.II | 1 | F8 | e |  | + |
| 839 | CheW.III | 1 | F9 | + | + | + |

<sup>a</sup>Outlined in Figure 1. The CheW.I *Architecture* splits into three *Classes*.<sup>36</sup>
<sup>b</sup> From <sup>36</sup>. The two CheW-like domains in CheA.V and CheA.VI *Architectures* belong to different *Classes*.
<sup>c</sup>Our sample does not contain enough representatives for analysis of system categories F3, F11-F17.
<sup>d</sup>Wuichet & Zhulin<sup>32</sup> noted CheA proteins modified only by C-terminal receiver domains (CheA.III or CheA.IV) were consistently observed in F3 chemotaxis systems. However, our PhyDCA scores in Dataset S4 do not support linkage of either the CheA.III or CheA.IV *Architectures* to either F3 or F4 chemotaxis systems. <sup>e</sup>Although the CheA.V, CheA.VII, CheA.VIII, CheA.X, and CheW.II *Architectures* contain additional domains with respect to canonical CheA or CheW *Architectures* and are sufficiently abundant to be included in <sup>36</sup>, these *Architectures* were apparently not observed sufficiently consistently in specific chemotaxis system categories to be noted by Wuichet & Zhulin<sup>32</sup>, who analyzed a much smaller sample size. <sup>f</sup>The subset of F7 chemotaxis systems that contain CheZ. See Figure 2. <sup>g</sup>Did not sort with a specific chemotaxis system category in Figure 2 and did not make sufficiently strong connections to be included in Figure 4.

The information presented by Figure 2 is challengingly dense, but salient features can be identified and discussed most easily using a grid coordinate system in which blocs are identified by assigned chemotaxis system category (rows, containing individual protein components; labeled on left with silver boxes) and representative proteome cluster (columns, containing distinct organisms; labeled on bottom with numbers as proteome clusters). Several individual rows in Figure 2 exhibited distributions from which we could glean additional insights. Prominent components spanning numerous proteome clusters included mcp.44H, mcp.24H, mcp.40H, mcp.34H, and mcp.36H (found in F1, F6, N/A, F8, and F7 blocs respectively); CheW.IB (F8 blocs); CheW.IA, which includes most CheW proteins (N/A blocs above F8); and CheA.I, the simplest and most common CheA *Architecture* (F7 blocs). The listed components are known to constitute the core signaling pathway shared by all chemotaxis systems. CheV.I (the sole version of CheV detected in any significant abundance; see ^36^), found in nearly a third of all chemotaxis systems in nature, also spanned many proteome clusters (F6 blocs). Strikingly, the occurrence of CheV.I correlated well with the mcp.40H type of chemoreceptor, strongly suggesting preferential interaction(s) (N/A blocs above F8).

Figure 2 also revealed several key features shared by the MAC1/2 chemotaxis categories. Methyl-accepting coiled-coil (MAC) proteins are closely related to chemotaxis proteins but, to the best of our knowledge, have not been experimentally characterized in any respect. MAC proteins include apparent chemoreceptor and kinase domains, and either incorporate (MAC1) or are associated with (MAC2) CheB and CheR related domains.^32^ It is unclear whether MAC proteins are evolutionary precursors of canonical chemotaxis systems or degenerate remnants. Proteome clusters 16, 17, and 18 featured high concentrations of organisms encoding MAC1 and/or MAC2 components but seemingly lacked other types of chemotaxis systems (MAC1 and MAC2 blocs). MAC systems were also scattered across numerous proteome clusters in Figure 2, rather than remaining constrained to contiguous blocs. Such a distribution suggested substantial phylogenetic prolificacy. Wuichet and Zhulin found that ∼80% of species with MAC systems encode additional chemotaxis systems.^32^ The distribution of MAC systems seen in Figure 3, based on our much larger sample size, supported and strengthened the original observation.

### 3.4 Co-evolutionary analysis reveals functional communities of chemotaxis components

Due to the complex nature of the information represented in Figure 2 (and the relative inability of the human eye to untangle multivariate correlations), we sought a way to simplify and quantify the co-occurrence probabilities of individual components observed in the various chemotaxis systems in nature. We used Phyletic Direct Coupling Analysis (PhyDCA) as described in Materials and Methods to quantify pairwise phyletic couplings between chemotaxis components and displayed the 4% strongest positive couplings (for which the presence of one component accurately predicts the presence of the other) in Figure 4.

The relationships revealed by Figure 4 largely corroborated the clustering patterns of Figure 2 and the assignments made in Table 2. Individual chemotaxis components typically associated closely with others of the same system category (represented by shared colors in Figure 4), but less strongly to nodes outside the same group. The network representation was particularly useful for visualizing relationships between the disparate categories/chemoreceptors and the CheW-/CheV-lineage *Architectures*. Most of the chemotaxis categories could be traced to a least one CheW- lineage and CheA-lineage component in relatively short order. Category F2 components and mcp.48H appeared as a cluster unconnected to other components in Figure 4, but all exhibited couplings in the top ∼10% to CheA.II (Dataset S4) and hence linked to the F1 chemotaxis system.

Figure 2 shows four groups of components (labeled N/A or uncategorized) that sorted into isolated blocs rather than associating with the standard chemotaxis system categories. Figure 4 suggests that the components were not associated with one another through co-evolutionary processes, but rather were dispersed and associated with a diverse range of other proteins. One explanation for the differences between the results in Figures 2 and 4 is correlation of individual components with multiple chemotaxis system categories. Such a phenomenon would likely facilitate linkage in Figure 4 but confound the clustering procedure used for Figure 2.

The strong connections between components observed in Figure 4 allowed us to confirm many of the chemotaxis system class assignments proposed in Table 2 (based on observations from Figure 2), as well as to infer several additional novel assignments. In particular, inspection of Figure 4 provides insights about how various *Architectures* and *Classes* of CheW and CheV scaffold proteins assort across the various chemotaxis system categories. Insights into the minor CheW.IA and CheW.IC *Architectures*, as well as CheV.I, can be found in the Discussion. Additional comments on CheA.IB, CheW.II, and CheW.III *Architectures* are included in Text S1.

### 3.5 Negative phyletic couplings reveal putative overlapping functionality among specific chemotaxis components

The PhyDCA model can also be used to predict negative phyletic couplings, i.e., the presence of one component in a proteome disfavors the presence of another.^47^ Logic suggests that components in such a scenario likely share overlapping (or at least closely related) functionalities (sometimes referred to as “alternative” solutions). For example, the top negative coupling within Dataset S4 involved the components cher.F8 and cher.Uncat, presumably because both methyltransferases serve highly similar functions. The third strongest negative coupling involved CheW.II and CheW.III. Curiously, few other instances of anticorrelated components with presumably similar functions are present in the list of top negative phyletic pairs, with most entries involving disparate component types (and are therefore not likely to be consequences of convergent evolution). The only exceptions (from the top 4% strongest anticorrelated phyletic pairs) were CheW.IA with CheW.IC (both CheW-lineage scaffolds), chec.F1 with chez.F7 (both phosphatases), CheA.I with CheA.II (the two most abundant CheA- lineage kinases), cher.F10 with cher.F5 (both methyltransferases), cheb.F1 with cheb.F10 (both methylesterases), cher.F1 with cher.F8 (both methyltransferases), cheb.F7 with cheb.F8 (both methylesterases), and finally cheb.F10 with cheb.F5 (both methylesterases).

## 4 DISCUSSION

### 4.1 Many combinations of chemotaxis system categories within a species are non-random

The observations described in this report, and the conclusions made or inferred, were only possible because we sampled proteomes from a large number of distinct organisms. The results of unsupervised two-dimensional sorting of species according to their content of chemotaxis system categories (Figures 2 and 3) strongly implies that the distribution of systems is substantially non-random. It is conceptually simple to sort species encoding only one category into a bloc (e.g., F1 bloc in Column 1 of Figure 2 or Column 6 of Figure 3). However, most species in our dataset encoded more than one chemotaxis system. Non-random combinations of systems would preclude two-dimensional sorting into blocs. The bloc data structure observed in Figures 2 and 3 suggested that a restricted subset of combinations of chemotaxis systems is evolutionarily favored. Preferred combinations must be either ancient (passed on to descendants of common ancestors) and/or are synergistically beneficial (arose independently multiple times). In either case, horizontal gene transfer of chemotaxis systems between species has not erased the pattern of combinatorial preferences in nature. Though members of some prokaryotic phyla tend to encode particular categories of chemotaxis systems, the topologies of the chemotaxis and species classification trees do not match^32, 51^, implying different evolutionary paths. The primary combinations of chemotaxis system categories (Table 1) and their non-random nature (Figures 2 and 3) may provide clues into the rise of various chemotaxis systems.

Figures 2 and 3 revealed large scale qualitative features of the data. Closer quantitative evaluation (Tables S1-S3) revealed more complex features. Many combinations of systems co-occurred more or less frequently than expected (i.e., a non-random pattern). Overall, our findings imply that most flagellar chemotaxis systems are not independent from one another. This would be expected if multiple systems control the same flagellar motors (e.g., having closely related CheY proteins to control the same motor could be advantageous). In contrast, ACF and Tfp systems appeared to act independently from each other, as did MAC1 and MAC2 systems.

### 4.2 Apparent functional specialization of proteins containing CheW-like domains

CheW-like domains primarily occur in CheA, CheV, and CheW proteins, but the CheA- and CheW-lineages exhibit substantial diversity (Figure 1).^36^ Figure 2 shows that CheA.I (the most abundant CheA *Architecture*), CheW.IA and CheW.IB (which together comprise almost all CheW proteins), CheV.I (the sole abundant CheV *Architecture*), and many types of MCPs were widely distributed across proteomes, implying an ability to function in multiple chemotaxis systems. However, our analyses (summarized in Table 2) strongly suggest that many *Architectures* of proteins containing CheW-like domains are associated with particular chemotaxis system categories, implying functional specialization. We confirmed four assignments made by Wuichet and Zhulin^32^ and made 17 new assignments. The additional assignments provide a rich foundation for future mechanistic investigations.

### 4.3 Insights into functions of uncommon Class 2 and Class 6 CheW-like domains

Most CheW-like domains in CheW proteins belong to *Class* 1 (Figure 1, Ref. ^36^). Co-occurrence data provide potential insight into proteins with CheW.I Architecture but less common *Class* 2 and *Class* 6 domains.

CheW.IA comprises *Class* 6 of CheW-like domains and makes up ∼20% of CheW.I proteins.^36^ CheW.IA is subtly distinguishable from CheW.IB and CheV.I CheW-like domains (*Class* 1) by some (but not all) methods of sequence analysis.^36^ In Figure 4, CheW.IA made direct connections to various chemoreceptors, but not to any CheA proteins (i.e., phyletic coupling of CheW.IA was stronger to MCPs than to CheA-lineage proteins). We speculate that the distinction between Class 1 and Class 6 CheW-like domains is that the latter exhibit greater specificity or preference for interactions with certain classes of MCPs (e.g., mcp.40H, mcp.38H). Note that CheW.IA sorted with mcp.40H in Figure 2 (N/A bloc above F8), but not with a specific chemotaxis system class. In a related observation, CheW.IA and CheW.IB account for nearly all single domain CheW proteins in nature. Both made strong direct connections to mcp.40H in Figure 4. CheW.IB also made direct connections with mcp.44H (strong) and mcp.36H (weaker), and CheW.IA made a direct connection (strong) with mcp.38H. Collectively, MCPs from 36H, 38H, 40H and 44H account for nearly 90% of all MCPs in nature (at least as encompassed by the RP35 dataset). When viewed in this way, the phyletic couplings between such prolific components makes sense. However, we again speculate that the “unique” interactions noted for CheW.IA and CheW.IB are rooted in preferences for distinct chemoreceptors. Although both likely share a robust ability to interact with mcp.40H, CheW.IA may also be able to interact with mcp.38H, whereas CheW.IB may be able to interact with both mcp.44H and mcp.36H.

CheW.IA was directly linked to F5, F6, and F7.z chemotaxis system categories in Figure 4. CheW.IB was directly linked to F1 and F7 components in Figure 4 and sorted with F8 in Figure 2. As the F7.z category diverged from F7 and became associated with CheA.VII and CheA.VIII rather than CheA.I *Architectures* (Table 2), CheW may have diverged in parallel from *Class* 1 CheW.IB (F7) to *Class* 6 CheW.1A (F7.z).

CheW.IC (*Class* 2) makes up ∼1% of CheW-like domains and shares characteristics of CheW-like domains found in both CheA and CheW proteins.^36^ CheW.IC did not sort with any known chemotaxis system category in Figure 2 (N/A bloc below F4), perhaps due to its low abundance in nature. The same rarity makes interpretation of the distribution of CheW.IC in Figure 2 challenging. However, CheW.IC was strongly linked with the MAC1 category in Figure 4 (corroborated upon close inspection of Figure 2). MAC systems incorporate both receptor and kinase functions into a single protein species, implying that they do not need CheW scaffold proteins to bridge the two elements. It is not known if any MAC proteins form arrays in conjunction with CheW proteins. In principle, arrays of MAC proteins could provide previously described advantages (e.g., sensitive signal detection, amplification, integration) of a canonical chemoreceptor array, but array properties might be constrained by the one-to-one relationship between intramolecular chemoreceptor and kinase functions in MAC proteins. Arrays could also facilitate adaptation, for example by allowing the CheB and/or CheR domains of MAC1 proteins to modify adjacent receptors, or the separate CheR proteins of MAC2 systems to localize to the array by molecular brachiation.^52^

### 4.4 CheW.II and CheW.III *Architectures* may interact with unusual MCPs

CheW.II and CheW.III *Architectures* are ∼1 to 3% as abundant as CheW.I *Architectures*.^36^ We sought to use negative PhyDCA pairings to infer the extent of overlapping roles between CheW proteins with various *Architectures*. Several results are consistent with the hypothesis that CheW.II and CheW.III are specialized to support interactions with particular MCPs. First, the third strongest negative PhyDCA coupling was between CheW.II and CheW.III (Dataset S4). A negative PhyDCA coupling suggests that CheW.II and CheW.III perform similar functions, so only one is needed. (An alternative possibility is that because CheW.II and CheW.III proteins are both rare, our sample size is too small to include species encoding both). Second, we are aware of only one experimental investigation of CheW.II or CheW.III function, which showed that CheW.III in *Vibrio cholerae* linked an unusual type of MCP to CheA in cytoplasmic double-layered arrays.^53^ Finally, none of the top negative PhyDCA pairings involving CheW.II or CheW.III featured any other CheW-lineage *Architecture*, implying, along with frequent co-occurrences with other CheW proteins in Figure 2, that the CheW.II and CheW.III *Architectures* are not “alternative” solutions for the more standard CheW-lineage components. In other words, CheW.II probably does not replace two distinct single-domain CheW proteins, but likely serves novel functionalities. However, an apparently distinct role is constrained by the fact that, like the abundant CheW.IB, CheW.II and CheW.III contain Class 1 CheW-like domains (Figure 1, Ref. ^36^).

### 4.5 A potential link between CheV and CheC/CheZ phosphatases

CheV.I clustered with category F6 chemotaxis systems in Figure 2 but also appeared coincident with the F1 category. Figure 4 confirmed connections between CheV.I and the F1 and F6 categories. Figure 2 also showed a strong correlation between CheV.I and mcp.40H, which was again confirmed by Figure 4. However, CheV.I did not show a direct connection to any specific CheA *Architecture*. Curiously, besides a link with mcp.40H, the only other strong direct correlations formed by CheV.I involved the phosphatases chez.F6/chec.F1 and the methyltransferase cher.F1. The role(s) of CheV-lineage proteins and their attached receiver domains are poorly understood. Some evidence suggests that CheV is involved in the chemotaxis adaptation process^31^, making the correlation between CheV.I and the CheC/CheR components of the F1 class^31^ understandable. However, the nature of the connection between CheV.I and chez.F6 is less clear and raises the concept of CheZ (and possibly the CheC of category F1) acting upon the attached phosphorylatable receiver domain of CheV.I in the capacity of a phosphatase. In fact, CheZ has phosphatase activity toward one of the three CheV proteins in *H. pylori*.^54^ It is not known whether CheZ distinguishes between different CheV proteins based on their CheW-like and/or receiver domains.

### 4.6 Limitations of PhyDCA

The PhyDCA approach used to generate Figure 4 has several disadvantages that must be considered. One is that the observed phyletic couplings do not necessarily correspond to direct biophysical interactions. Strong coupling may also represent events such as genomic co-localization (a limitation the original authors circumvented by including an additional residue-level covariance analysis to predict likely direct interaction partners).^47^ Because of the narrow perspective of our study (i.e., focusing on chemotaxis systems, rather than entire proteomes), we believe that this disadvantage was minimized. However, as can be seen in Figure 2, paralogous chemotaxis components are common in bacteria, particularly for chemoreceptors. Introducing a residue-level analysis step may facilitate the untangling of specific paralog interactions,^47, 55^ especially matching the various CheW-like domains in CheA-, CheW-, and CheV-lineage proteins to their partner MCP components. Such an analysis would provide additional insight into the diverse chemotaxis systems found in nature. Additionally, PhyDCA (and many other phylogenetic profiling methods) relies on a binary presence/absence data structure to simplify data processing and interpretation. Excluding the substantial amount of paralogous protein data found in our microbial dataset very likely ignores valuable co-occurrence information. However, our two-pronged approach to the problem (using the full co-occurrence matrix to identify functional blocs/clusters in Figure 2 and using the transformed binary profile to create the simplified network representation in Figure 4) likely mitigates the issue.

### 4.7 General insights into occurrence and organization of prokaryotic chemotaxis systems

CheW-like domains play a central role in the signal transduction systems that regulate prokaryotic chemotaxis by linking receptors and kinases into large arrays. Almost all CheW-like domains occur in a limited number of *Architectures* of CheA-, CheW-, and CheV-lineage proteins (Figure 1).^36^ Furthermore, CheW-like domains have evolved into distinct functional *Classes* (Figure 1).^36^ We surveyed chemotaxis proteins encoded by nearly 1900 species (Dataset S1) and examined their distribution by both chemotaxis system category^32^ and by species/proteome. Successful unsupervised clustering of components into blocs (Figure 2) strongly suggested that the components were linked in both dimensions, leading to two central conclusions. First, many combinations of chemotaxis systems encoded by individual species (Table 1, Dataset S2) were non-random (Figure 3, Tables S1-S3). Specific co-occurrence patterns and frequencies (Dataset S3) should provide insights into evolution of chemotaxis systems. Second, we inferred probable functional associations between each *Architecture* of CheA-, CheW-, and CheV-lineage proteins and specific categories of chemotaxis systems (Figure 4, Table 2, Dataset S4). These assignments lay a foundation for future investigations into the mechanisms that underly apparent functional specialization of different chemotaxis protein *Architectures*.

## ACKNOWLEDGEMENTS

We thank Emily N. Kennedy and Sarah A. Barr for their insightful input and helpful discussions.

This work was funded by National Institutes of Health grant GM050860 to Robert B. Bourret. The content is solely the responsibility of the authors and does not necessarily represent the official views of the National Institute of General Medical Sciences or the National Institutes of Health.

