## Supplementary material for "Analysis of CheW-like domains provides insights into organization of prokaryotic chemotaxis systems": Text S1, Figure S1, Tables S1-S3

Revised for *Proteins: Structure, Function, and Bioinformatics*

5/31/22

Running Title: CheW-like domains and chemotaxis systems

*Corresponding author

### SUPPORTING INFORMATION

#### Text S1. Insights into CheW.IB, CheW.II, and CheW.III derived from Figure 4

CheW.IB (the most common form of CheW) sorted with chemotaxis system category F8 in Figure 2 but connected primarily with the F1 and F7 systems in Figure 4. This pattern is consistent with the fact that F1, F7, and F8 are the most abundant chemotaxis system categories (Dataset S3, Ref. [1]). CheW.IB linked with CheA.I and CheA.II (the two most common CheA *Architectures*[2]) in Figure 4. CheW.IB also connected to multiple types of MCPs in Figure 4. All described linkages are consistent with common core components utilized by many different categories of chemotaxis systems.

A recent report noted that the F7 system of *E. coli* likely evolved from merging ancient versions of the F6 and F7 systems.[3] Wuichet and Zhulin were unable to assign a characteristic chemoreceptor to the F6 system category.[1] Whereas the heatmap in Figure 2 implies that the appropriate MCP may be mcp.24H, the phyletic coupling network suggests that it could also be mcp.40H, via an indirect (but relatively strong) correlation with CheW.IA. Similarly, category F7 is strongly associated with mcp.36H (as previously reported[1]) but may also be linked to mcp.40H via CheW.IB.

CheW.II sorted with ACF systems in Figure 2 but made its strongest connection to category F8 systems in Figure 4. In contrast to most other types of CheW proteins, CheW.II did not make strong direct connections to individual MCPs in Figure 4, implying that CheW.II proteins may be promiscuous and interact with multiple different types of chemoreceptors. An alternative interpretation equally consistent with the data is that CheW.II proteins do not interact with MCPs at all but have an as yet to be determined function. We are not aware of any experimental investigation of CheW.II proteins. CheA.IX did not sort with a specific chemotaxis system category in Figure 2 (N/A bloc below F4), but connected to CheW.II in Figure 4, suggesting an association with the category F8 and ACF systems.

Similarly, CheW.III (category F9), CheA.XI (unassigned in Figure 2, N/A bloc below F4), and CheA.XII (category F4) were strongly interconnected in Figure 4, suggesting CheA.XI may belong to both the class F4 and F9 chemotaxis systems. Belonging to multiple categories of chemotaxis systems could explain a failure to sort coherently in Figure 2. There also may be functional reasons for these apparent interconnections. The CheA.IX *Architecture* lacks Hpt domains with phosphorylation sites, whereas the CheA.XI and CheA.XII *Architectures* lack the catalytic and ATP-binding HATPase_c domain (Figure 1). Therefore, all three must interact with other CheA proteins to participate in phosphotransfer reactions.

**Figure S1. Row dendogram for Figure 2.**

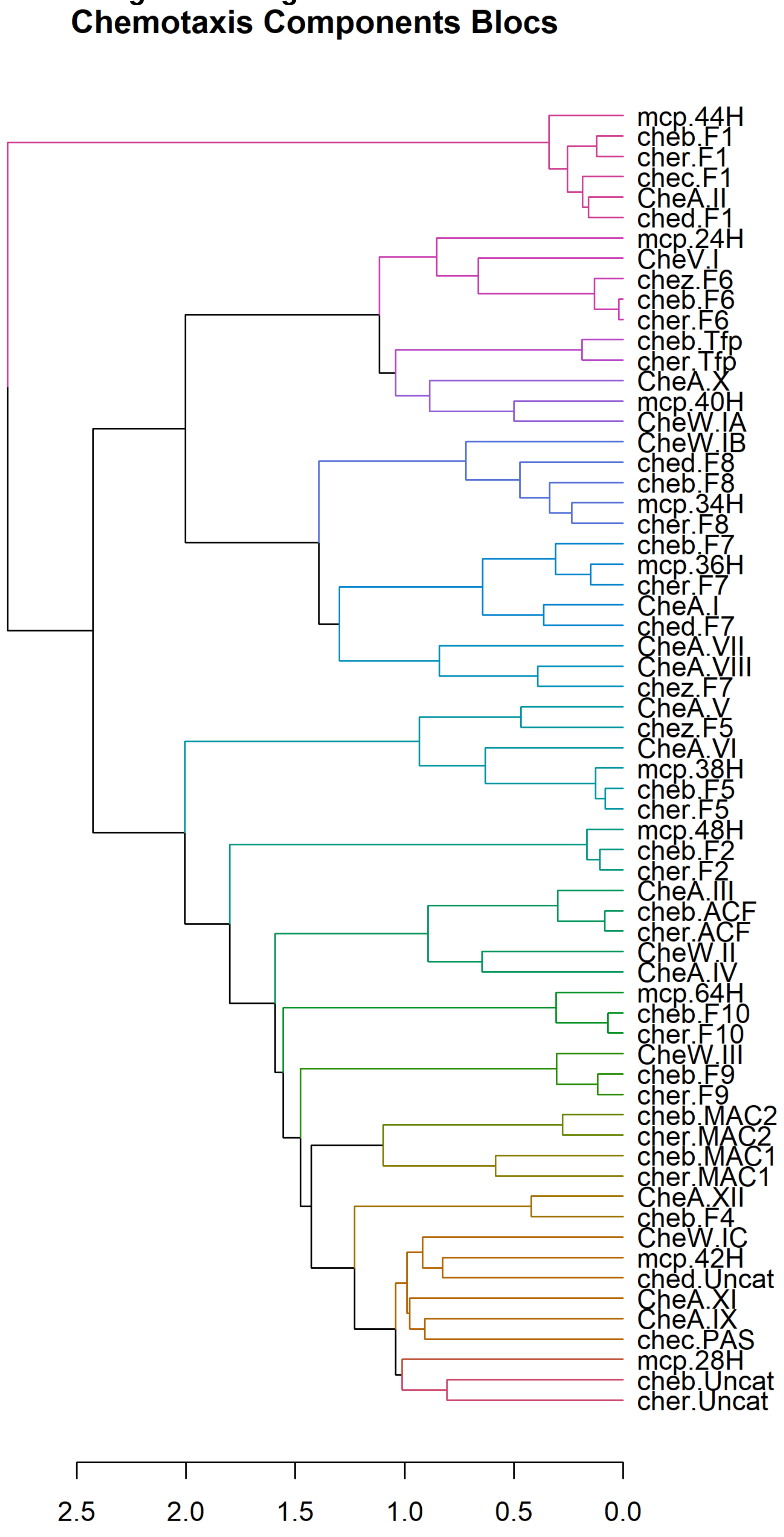

**Table S1. Expected and observed frequencies of pairwise combinations of chemotaxis system categories within branches of phylogenetic tree**

| | | | | Both System 1 and System 2 | | |
|---|---|---|---|---|---|---|
| System 1 | Frequency[a] | System 2 | Frequency | Number Observed | Frequency Expected[b] | Frequency Observed[c] |
| *Branch 2A*[d] | | | | | | |
| F1 | 0.45 | F2 | 0.017 | 28 | 0.0077 | 0.016 |
| *Branch 2C* | | | | | | |
| F5 | 0.17 | F6 | 0.091 | 6 | 0.015 | 0.0033 |
| *Branch 2B* | | | | | | |
| F4 | 0.012 | F9 | 0.062 | 2 | 0.00074 | 0.0011 |
| F4 | 0.012 | F10 | 0.013 | 1 | 0.00016 | 0.00056 |
| F9 | 0.062 | F10 | 0.013 | 8 | 0.00081 | 0.0045 |
| *Branch 1* | | | | | | |
| F7.z | 0.090 | F7 | 0.38 | 158 | 0.035 | 0.088 |
| F7.z | 0.090 | F8 | 0.20 | 112 | 0.018 | 0.062 |
| F7 | 0.38 | F8 | 0.20 | 277 | 0.076 | 0.15 |
| *Branch 3* | | | | | | |
| ACF | 0.13 | Tfp | 0.039 | 7 | 0.0051 | 0.0039 |
| *"Branch 4"* | | | | | | |
| MAC1 | 0.29 | MAC2 | 0.19 | 142 | 0.055 | 0.079 |

[a]Fraction of genomes containing indicated system from Dataset S3.
[b]Expected Frequency = (Frequency of System 1)(Frequency of System 2)
[c]Red indicates observed ≤ 0.5x expected, yellow indicates observed within 2x of expected, green indicates observed ≥ 2x expected
[d]The phylogenetic tree in Figure 7 of Wuichet & Zhulin[1] that forms the basis of classification of chemotaxis system categories has three main branches, arbitrarily numbered here. Branch 2 has three main sub-branches (A, B, C).

**Table S2. Expected and observed frequencies of pairwise combinations of chemotaxis system categories between branches of phylogenetic tree[a]**

| | | | | Both System 1 and System 2 | | |
|---|---|---|---|---|---|---|
| System 1 | Frequency[b] | System 2 | Frequency | Number Observed | Frequency Expected[c] | Frequency Observed[d] |
| *Branches 2A and 2C[e]* | | | | | | |
| F1 | 0.45 | F5 | 0.17 | 101 | 0.077 | 0.056 |
| F1 | 0.45 | F6 | 0.091 | 48 | 0.041 | 0.027 |
| *Branches 2A and 2B* | | | | | | |
| F1 | 0.45 | F9 | 0.062 | 52 | 0.028 | 0.029 |
| *Branches 2A and 1* | | | | | | |
| F1 | 0.45 | F7.z | 0.090 | 1 | 0.041 | 0.00056 |
| F1 | 0.45 | F7 | 0.38 | 236 | 0.17 | 0.13 |
| F1 | 0.45 | F8 | 0.20 | 122 | 0.090 | 0.068 |
| *Branches 2C and 2B* | | | | | | |
| F5 | 0.17 | F9 | 0.062 | 40 | 0.011 | 0.022 |
| F6 | 0.091 | F9 | 0.062 | 10 | 0.0056 | 0.0056 |
| *Branches 2C and 1* | | | | | | |
| F5 | 0.17 | F7.z | 0.090 | 0 | 0.015 | < 0.00056 |
| F5 | 0.17 | F7 | 0.38 | 94 | 0.065 | 0.052 |
| F5 | 0.17 | F8 | 0.20 | 73 | 0.034 | 0.041 |
| F6 | 0.091 | F7.z | 0.090 | 0 | 0.0082 | < 0.00056 |
| F6 | 0.091 | F7 | 0.38 | 95 | 0.035 | 0.053 |
| F6 | 0.091 | F8 | 0.20 | 49 | 0.018 | 0.027 |
| *Branches 2B and 1* | | | | | | |
| F9 | 0.062 | F7.z | 0.090 | 12 | 0.0056 | 0.0067 |
| F9 | 0.062 | F7 | 0.38 | 51 | 0.024 | 0.028 |
| F9 | 0.062 | F8 | 0.20 | 47 | 0.012 | 0.026 |
| *Branches 3 and 4* | | | | | | |
| ACF | 0.13 | MAC1 | 0.29 | 115 | 0.038 | 0.064 |
| ACF | 0.13 | MAC2 | 0.19 | 56 | 0.025 | 0.031 |
| Tfp | 0.039 | MAC1 | 0.29 | 13 | 0.011 | 0.0072 |
| Tfp | 0.039 | MAC2 | 0.19 | 13 | 0.0074 | 0.0072 |

[a]Does not include 20 combinations involving minor classes F2, F4, and F10, which are too infrequent to yield testable frequency predictions.

[b]Fraction of genomes containing indicated system from Dataset S3.

[c]Expected Frequency = (Frequency of System 1)(Frequency of System 2)

[d]Red indicates observed ≤ 0.5x expected, yellow indicates observed within 2x of expected, green indicates observed ≥ 2x expected

[e]The phylogenetic tree in Figure 7 of Wuichet & Zhulin[1] that forms the basis of classification of chemotaxis system categories has three main branches, arbitrarily numbered here.  Branch 2 has three main sub-branches (A, B, C).

**Table S3. Expected and observed frequencies of pairwise combinations of flagellar chemotaxis system categories with ACF, Tfp, MAC1 or MAC2 systems**

| | | | | Both System 1 and System 2 | | |
|---|---|---|---|---|---|---|
| System 1 | Frequency[a] | System 2 | Frequency | Number Observed | Frequency Expected[b] | Frequency Observed[c] |
| *Branch 2A[d]* | | | | | | |
| F1 | 0.45 | ACF | 0.13 | 23 | 0.059 | 0.013 |
| F1 | 0.45 | Tfp | 0.039 | 2 | 0.018 | 0.0011 |
| F1 | 0.45 | MAC1 | 0.29 | 116 | 0.13 | 0.065 |
| F1 | 0.45 | MAC2 | 0.19 | 129 | 0.086 | 0.072 |
| F2 | 0.017 | ACF | 0.13 | 4 | 0.0022 | 0.0022 |
| F2 | 0.017 | Tfp | 0.039 | 0 | 0.00066 | <0.00056 |
| F2 | 0.017 | MAC1 | 0.29 | 3 | 0.0049 | 0.0017 |
| F2 | 0.017 | MAC2 | 0.19 | 1 | 0.0032 | 0.00056 |
| *Branch 2C* | | | | | | |
| F5 | 0.17 | ACF | 0.13 | 49 | 0.022 | 0.027 |
| F5 | 0.17 | Tfp | 0.039 | 0 | 0.0066 | <0.00056 |
| F5 | 0.17 | MAC1 | 0.29 | 100 | 0.049 | 0.056 |
| F5 | 0.17 | MAC2 | 0.19 | 43 | 0.032 | 0.024 |
| F6 | 0.091 | ACF | 0.13 | 26 | 0.012 | 0.014 |
| F6 | 0.091 | Tfp | 0.039 | 42 | 0.0035 | 0.023 |
| F6 | 0.091 | MAC1 | 0.29 | 42 | 0.026 | 0.023 |
| F6 | 0.091 | MAC2 | 0.19 | 28 | 0.017 | 0.016 |
| *Branch 2B* | | | | | | |
| F4 | 0.012 | ACF | 0.13 | 5 | 0.0016 | 0.0029 |
| F4 | 0.012 | Tfp | 0.039 | 0 | 0.00047 | <0.00056 |
| F4 | 0.012 | MAC1 | 0.29 | 9 | 0.0035 | 0.0050 |
| F4 | 0.012 | MAC2 | 0.19 | 3 | 0.0023 | 0.0017 |
| F9 | 0.062 | ACF | 0.13 | 34 | 0.0081 | 0.019 |
| F9 | 0.062 | Tfp | 0.039 | 2 | 0.0024 | 0.0011 |
| F9 | 0.062 | MAC1 | 0.29 | 51 | 0.018 | 0.028 |
| F9 | 0.062 | MAC2 | 0.19 | 21 | 0.012 | 0.012 |
| F10 | 0.013 | ACF | 0.13 | 14 | 0.0017 | 0.0078 |
| F10 | 0.013 | Tfp | 0.039 | 0 | 0.00051 | <0.00056 |
| F10 | 0.013 | MAC1 | 0.29 | 16 | 0.0038 | 0.0089 |
| F10 | 0.013 | MAC2 | 0.19 | 7 | 0.0025 | 0.0039 |
| *Branch 1* | | | | | | |
| F7.z | 0.090 | ACF | 0.13 | 31 | 0.012 | 0.017 |
| F7.z | 0.090 | Tfp | 0.039 | 2 | 0.0035 | 0.0011 |

| | | | | | | |
|---|---|---|---|---|---|---|
| F7.z | 0.090 | MAC1 | 0.29 | 49 | 0.026 | 0.027 |
| F7.z | 0.090 | MAC2 | 0.19 | 32 | 0.017 | 0.018 |
| F7 | 0.38 | ACF | 0.13 | 86 | 0.049 | 0.048 |
| F7 | 0.38 | Tfp | 0.039 | 33 | 0.015 | 0.018 |
| F7 | 0.38 | MAC1 | 0.29 | 93 | 0.11 | 0.052 |
| F7 | 0.38 | MAC2 | 0.19 | 111 | 0.072 | 0.062 |
| F8 | 0.20 | ACF | 0.13 | 38 | 0.026 | 0.021 |
| F8 | 0.20 | Tfp | 0.039 | 15 | 0.0078 | 0.0083 |
| F8 | 0.20 | MAC1 | 0.29 | 153 | 0.058 | 0.085 |
| F8 | 0.20 | MAC2 | 0.19 | 24 | 0.038 | 0.013 |

[a]Fraction of genomes containing indicated system from Dataset S3.

[b]Expected Frequency = (Frequency of System 1)(Frequency of System 2)

[c]Red indicates observed ≤ 0.5x expected, yellow indicates observed within 2x of expected, green indicates observed ≥ 2x expected

[d]The phylogenetic tree in Figure 7 of Wuichet & Zhulin[1] that forms the basis of classification of chemotaxis system categories has three main branches, arbitrarily numbered here.  Branch 2 has three main sub-branches (A, B, C).

Provided as separate .xlsx files:

**Dataset S1. Component Frequency Matrix.** Lists the number of each of 64 different types of chemotaxis system components encoded by each of ~1900 prokaryotic genomes. Components from relatively infrequent chemotaxis system categories F3 and F11 through F17 are not included.

**Dataset S2. Che System Category Combinations.** Presence/absence combinations of common chemotaxis system categories (excluding F3 and F11 through F17) encoded by inventoried genomes.

**Dataset S3. Che System Category Frequencies**. Frequencies of chemotaxis system categories observed among the genomes analyzed in this work and by Wuichet & Zhulin 2010.

**Dataset S4. PhyDCA Couplings.** 3,486 pairwise couplings between all 84 types of chemotaxis system components, including the infrequent F3 and F11 through F17 categories.
